# Addressing Underestimation of Waterborne Disease Risks Due to Fecal Indicator Bacteria Bound in Aggregates

**DOI:** 10.1101/2024.07.31.605961

**Authors:** Dan E. Angelescu, David Abi-Saab, Raphael Ganaye, David Wanless, Joyce Wong

## Abstract

Microbiological water quality monitoring is critical for managing waterborne disease risk; currently, regulations rely on quantifying culturable fecal indicator bacteria using traditional culture-based methods. These approaches cannot distinguish between planktonic forms and aggregates harboring higher loads of bacteria and associated pathogens, potentially underestimating exposure risks. By using size fractionation and ALERT, an automated rapid method for comprehensive quantification of culturable bacteria, we reveal widespread and substantial presence of aggregate-bound indicator bacteria across a variety of water matrices and geographies. We observe comprehensive bacteria counts exceeding traditional method counts by significant multiples (e.g., 3.4× on average at the Seine River 2024 Olympic venue, occasionally 100× in irrigation canals and wastewater plant effluent). The results, corroborated by microscopic and molecular analyses, highlight a major systematic bias in global water safety regulatory frameworks. Automated comprehensive culture-based rapid quantification methods could provide higher-accuracy risk assessments, enabling effective monitoring, including in remote and resource-limited settings.

## 1. Introduction

Waterborne pathogens of fecal origin pose a significant global health risk, primarily transmitted through contaminated drinking water, food, and recreational exposure. Microbiological contamination is responsible for over 1.5 million annual diarrheal fatalities^1^ and ranks as the fourth leading cause of mortality in young children globally^2^, with risk burden being disproportionately borne by sub-Saharan Africa and parts of Asia, where reliance on unsafe water sources persists^1^. In contrast, developed nations have seen a decrease in waterborne diseases, thanks to improved infrastructure and rigorous water safety protocols, including quantitative microbial risk assessments^3^. Even there, economic repercussions of unsafe water remain nevertheless substantial, with annual costs of billions of dollars in North America alone being attributed to *E. coli* outbreaks associated with leafy greens consumption^4,5^ and to illnesses from recreational water activities^6^. The climate crisis and associated extreme weather patterns exacerbate these issues by compromising water quality and quantity^7^ and prompting a pivot towards alternative sources like recycled wastewater^8^, which carries its own risks. Recreational use of surface waters, including swimming, is acknowledged for significant psychological and wellness benefits^9^ and has witnessed a surge in popularity since the COVID-19 lockdowns. This trend further underscores the importance of effective water quality monitoring—a factor highlighted by event cancellations, such as those in France during the 2023 preparatory season for the 2024 Paris Olympics, due to *E. coli* pollution from heavy rainfall and infrastructure issues^10^, or recurrent illness outbreaks at triathlon events in the UK despite testing results deemed acceptable^11^. These incidents highlight the ongoing challenges in securing recreational water safety.

Epidemiological research correlating the prevalence of viable and culturable fecal indicator bacteria (FIB), such as *E. coli* and enterococci, with the incidence of waterborne disease^12,13,14^ has led to their widespread use in water quality regulations. *E. coli* is the most common indicator for assessing the microbiological quality of potable water, inland surface water, and treated municipal wastewater quality, whereas seawater regulations tend to rely on enterococci, sometimes complemented by *E. coli*. In the developed world, drinking water assessments rely primarily on *E. coli* presence/absence tests. In resource-constrained settings, by contrast, quantitative *E. coli* measurements are essential for prioritizing among available drinking water sources^15^. Surveys show that exposure to fecal contamination through drinking water is widespread^1^, with over 80% of the population in some African countries only having access to untreated water containing various levels of fecal contamination^16^. While traditional FIB quantification techniques using grab sampling^17,18,19^ combined with culture-based methods — such as Membrane Filtration (MF)^20,21,22,23^ and plating, or Most Probable Number (MPN) ^24,25,26,27^ — are ubiquitously used in regulatory microbiological water analysis, they fail to distinguish between free-floating (planktonic) and aggregate-bound bacteria^28,29^. These limitations of traditional FIB quantification techniques can lead to skewed public health risk assessments in presence of FIB aggregates. Current methodologies do not include homogenization procedures more forceful than sample agitation^17–27,30^, which lacks the shear stresses necessary to mechanically disperse aggregates. While vigorous homogenization strategies such as sonication or bead beating are incompatible with culture-based methods due to the concurrent lysing of live bacteria, comprehensive glucuronidase activity assays have indicated the presence of significant amounts of particle-attached FIB in rivers^31^, particularly following combined sewer overflows (CSOs)^32^.

Recent advancements in water quality analysis include the development of culture-independent molecular assays for the detection and quantification of FIB. These assays incorporate thorough homogenization steps, such as bead-beating, nucleic acid extraction and concentration, followed by quantification via quantitative polymerase chain reaction (qPCR) or droplet digital PCR (ddPCR) methods, which include a reverse transcription (RT) step for RNA (RT-qPCR, RT-ddPCR). RT assays are typically tailored to target viable FIB^33^, whereas conventional molecular assays amplify DNA from all cells, necessitating additional measures to inhibit amplification from non-viable cells^34^ when assessing live FIB populations^35^. While faster than culture-based methods, molecular methods are particularly complex, requiring well-equipped molecular biology laboratories and highly trained personnel, which limits their widespread application, particularly in remote or resource-constrained settings where water quality challenges and public health impacts are often most acute.

In the United States the Environmental Protection Agency has endorsed certain molecular methods for the quantification of specific FIB, such as enterococci^36,37^. However, broader adoption of these methods is hindered by the difficulty of correlating qPCR results with those from MF or MPN techniques, and by a divergent response to standard disinfection treatments, with qPCR not exhibiting the anticipated reductions^38^. Nevertheless, epidemiological studies employing qPCR have demonstrated its potential as a reliable indicator of gastrointestinal disease risk^39^, particularly in children^40^. Often qPCR yields more consistent results than culture-based methods^41^, which are prone to produce variable risk assessments and underestimate the potential risk^42^, especially at lower FIB levels^43^.

Currently, despite recent estimates that 40-80% of all cells overall are bound within various types of aggregates and biofilms^44^, including the ubiquitous presence of unattached bacterial aggregates^45^ in freshwater, marine, and wastewater environments^46^, relatively little research exists in relation to their association with FIB or pathogens. FIB aggregates have been documented in stormwater and wastewater,^46,47,48^ yet their contribution to the known variability^49,50^ and apparent inconsistencies in FIB data and the implications for epidemiological risk have not been established, particularly against the backdrop of the intrinsic limitations of traditional methods to accurately count aggregate-bound FIB.

Here we investigate the partitioning of *E. coli* bacteria between planktonic and bound states on aggregates of different sizes, applying a novel methodology based on combining filtration-based size fractionation with simultaneous *E. coli* assessments using the traditional MPN and MF methods and ALERT, a rapid microbiological method^51^ providing whole-sample comprehensive quantification of *E. coli* regardless of their free or aggregate-bound state. We observe presence of aggregate-bound *E. coli* bacteria in most samples, in quantities often exceeding the planktonic load by significant multiples and in certain cases by over two orders of magnitude. Our measurements are corroborated by microscopic confirmation of aggregate-bound bacteria, complemented by molecular analyses of *E. coli* RNA. Our results demonstrate that, due to their insensitivity to aggregate-bound FIB, traditional MPN and MF methods may be routinely underestimating the overall FIB load, often by significant yet highly variable factors. Such methodological limitations can result in highly inaccurate risk assessments, that fail to fully capture the potential for waterborne disease transmission. The observed variability prevents simple corrective action such as readjusting the risk thresholds, implying that new risk assessment approaches need to be developed that are based on measuring the comprehensive FIB load, including bacteria present on aggregates.

## 2. Methods

### 2.1 Water sampling

Samples were collected using standard grab sampling techniques except where otherwise noted, and then either processed on-site (Brighton swimming competition samples) or transported to the laboratory in a cooler with ice. In the laboratory, samples were either processed immediately (Seine River samples, Los Angeles River samples), or stored in the fridge and processed later, within recommended holding times. Seine River sampling was performed near the Alexandre III bridge (GPS coordinates: 48.863248, 2.314508). Treated municipal wastewater effluent sampling for ALERT measurements was performed automatically, using an ALERT System V2 (Manufacturer: Fluidion).

### 2.2 Sample filtration and size fractionation

Filtration at 5 µm for the Seine River and Los Angeles River samples was performed using a sterilized 47 mm VWR membrane filtration apparatus (borosilicate glass) coupled to a laboratory vacuum pump (KNF), and 47 mm-diameter hydrophilic 5 µm cellulose nitrate membrane filters (Whatman). Filtration for all the other studies was performed by pushing the sample manually, using a sterile 30 mL syringe, through a 5 µm cellulose nitrate Luer Lock syringe filter. Filtration for the treated municipal wastewater effluent sampling was performed automatically, by coupling 5 µm sterile glass microfiber filters to the ALERT System V2 cartridges. The same procedure as for the 5 µm sample filtration was used for the size fractionation study, but first using a 47 mm-diameter hydrophilic 10 µm Nylon net membrane filter (Millipore). The 10 µm filtrate was then filtered again using the 5 µm filtration methodology.

### 2.3 MPN *E. coli* measurements

The IDEXX Colilert Quantitray-2000 system was used to perform all MPN measurements, following standardized protocols^24^. For environmental samples, typical sample dilutions of 1:10 to 1:100 were applied, using sterilized glassware and autoclave-sterilized de-ionized water (Millipore Elix® Essential 3 Water Purification System). The measurement values and confidence intervals were obtained from the IDEXX Quanti-Tray®/2000 MPN Table (per 100 mL) with 95% Confidence Limits, available online.

### 2.4 RT-qPCR *E. coli* measurements

All RT-qPCR FIB measurements were performed by Suez Rivages Pro Tech, using the GenSpot methodology, following published protocols^33^. The results were reported in cell-equivalent counts per 100mL.

### 2.5 ALERT *E. coli* measurements

ALERT measurements were performed using ALERT LAB portable *E. coli* analyzers (https://fluidion.com/products/analyzers/alert-lab) for all experiments, except the treated municipal wastewater effluent sampling, in which case an ALERT System V2 in-situ system was used instead (https://fluidion.com/products/analyzers/alert-v2). In all cases, the bioreagent containing the growth medium and 4-methylumbelliferyl-b-D-glucuronide (MUG) enzymatic substrate was used in liquid form, provided by the manufacturer either in a sterile syringe (ALERT LAB) or already loaded within a disposable measurement cartridge (ALERT System V2). For ALERT LAB, the raw or 5 µm-filtered sample (25 mL) was added together with 1 mL of bioreagent to a sterile vial and was briefly stirred, and then the measurement was started. For ALERT System V2, the system containing seven cartridges was attached to a tripod and installed through a trap in the effluent channel. Alternating cartridges had 5 µm in-line filters installed, and two samples were collected and analyzed in parallel (unfiltered and 5 µm-filtered). In all cases the measurement results and PDF data reports, including quality control data (raw signal curves, incubation temperature log, water temperature log), were downloaded from the Fluidion Data Analytics interface (https://data.secure-fluidion.com).

### 2.6 Bacterial direct staining procedures

Buffer preparation: 0.15 M NaCl buffer was prepared using high purity salt and autoclave-sterilized de-ionized water (Millipore Elix® Essential 3 Water Purification System). Ultrafiltration: A volume of 100 mL of sample was filtered using a sterilized 47 mm VWR membrane filtration apparatus (borosilicate glass) coupled to an KNF laboratory vacuum pump, and a 47 mm-diameter hydrophilic 0.2 µm black Nucleopore track-etched polycarbonate membrane filter (Whatman). BactoView^TM^ staining: A volume of 200 µl of 1X BactoView^TM^ Live Green solution (Biotium, REF: 40102-T) was prepared in buffer and was pipetted over the filter without removing it from the filtration apparatus, ensuring complete filter coverage. The filtration apparatus was then stored in the dark for 30 minutes. 100 mL of buffer were then passed through the filter to wash the unattached dye. WGA and DAPI staining: Volumes of 200 µl of 1X CF^®^488A WGA (Biotium, REF: 29022-1) and 1X DAPI (Biotium, REF: 40043) solutions were prepared in buffer. After sample filtration, the WGA solution was pipetted over the filter, ensuring complete coverage, after which the filtration apparatus was stored in the dark for 30 minutes. 100 mL of buffer were then passed through the filter to wash the unattached dye. The DAPI solution was then pipetted over the filter, ensuring complete coverage, after which the filtration apparatus was stored in the dark for 10 minutes. Finally, 50 mL of buffer were then passed through the filter to wash the unattached dye. All processing was done at room temperature, and without removing the filter from the apparatus.

### 2.7 Epifluorescence microscopy of stained bacteria

Microscopy setup: A Leica DMI 3000M inverted fluorescence microscope was used for all imaging. It was outfitted with an oil immersion objective (Leica N PLAN 50x/0,90 OIL), a fiber-coupled mercury lamp, and the A and I3 fluorescence dichroic filter cubes that allowed differential imaging of the DAPI and WGA stains. Sample preparation: The Nucleopore filter containing the dyed sample was carefully removed from the filtration apparatus using round-tipped sterilized metal tweezers. It was then cut in approximately 1 cm^2^ portions using scissors. A portion of the filter was then placed over a glass slide and secured to it using a small drop of N-type immersion oil. Additional drops of N-type immersion oil were placed over the filter and on the objective, respectively. The glass slide was then placed on the microscope stage. Imaging: Image recording was performed using a FHD V2.0 C-mount color camera, and images were saved to an SD memory card for further processing (background subtraction, noise reduction, contrast enhancement) using the Fiji software package.

## 3. Results

Our study, involving the analysis of 264 samples collected across four countries and from a wide variety of aquatic environments, principally focuses on investigating the prevalence of aggregate-bound FIB in the Seine River in Paris during the 2023 summer and fall seasons, at the anticipated 2024 Olympic venue. To this end we combine MPN and ALERT measurements to quantify the presence of aggregate-bound FIB, using a 5 µm filtration-based size threshold between planktonic and aggregate-bound *E. coli* bacteria. We analyze the consequences in terms of risk assessment discrepancies, according to international recreational norms. A size-fractionation is implemented to study the prevalence of FIB across aggregates of different sizes. Presence of aggregate-bound Gram-positive and Gram-negative bacteria is observed microscopically and the prevalence of aggregate-bound FIB is further confirmed using molecular RT-qPCR measurements. In the final part of the study, we extend the scope to a wide range of water matrices and locations, uncovering quasi-ubiquitous presence of aggregate-bound FIB in irrigation channels, inland and marine recreational waters, as well as in stormwater and treated wastewater intended for agricultural reuse.

### 3.1 Aggregated FIB observed at 2024 Olympic venue

From July to November 2023, we gathered time-series FIB concentration data (51 samples) at Pont Alexandre III on the Seine River, the projected site for the open-water events of the 2024 Paris Olympics. We utilized MPN (Colilert Quantitray-2000) and ALERT methods to quantify *E. coli* levels in both raw samples and those filtered through a 5 µm pore size, which excludes larger aggregates but allows passage of planktonic FIB and very small aggregates. A positive correlation between *E. coli* counts by all methods and precipitation was observed (Figure 1a), with expected concentration spikes correlated with rain events, due to CSOs, and an expected decline during drier conditions (*e.g.,* early to mid-September). All methods showed variations in FIB concentration of more than two orders of magnitude, with observed ranges (per 100mL) of 187 to 27,550 (for MPN), 110 to 19,400 (for 5 µm-filtered ALERT) and 314 to 42,500 (for unfiltered ALERT). The four highest values recorded by all methods were registered on October 26, October 24, August 2 and September 21, in the order of decreasing FIB load. The Open Water Swimming World Cup event scheduled on August 5-6 had to be canceled in the aftermath of the August 2 CSO episode, and despite relatively dry weather between mid-August and early September, elevated *E. coli* levels were observed and led to the partial cancellation of a World Triathlon event. The high FIB levels were later attributed to a valve malfunction on the combined sewer network.

**Figure 1.**
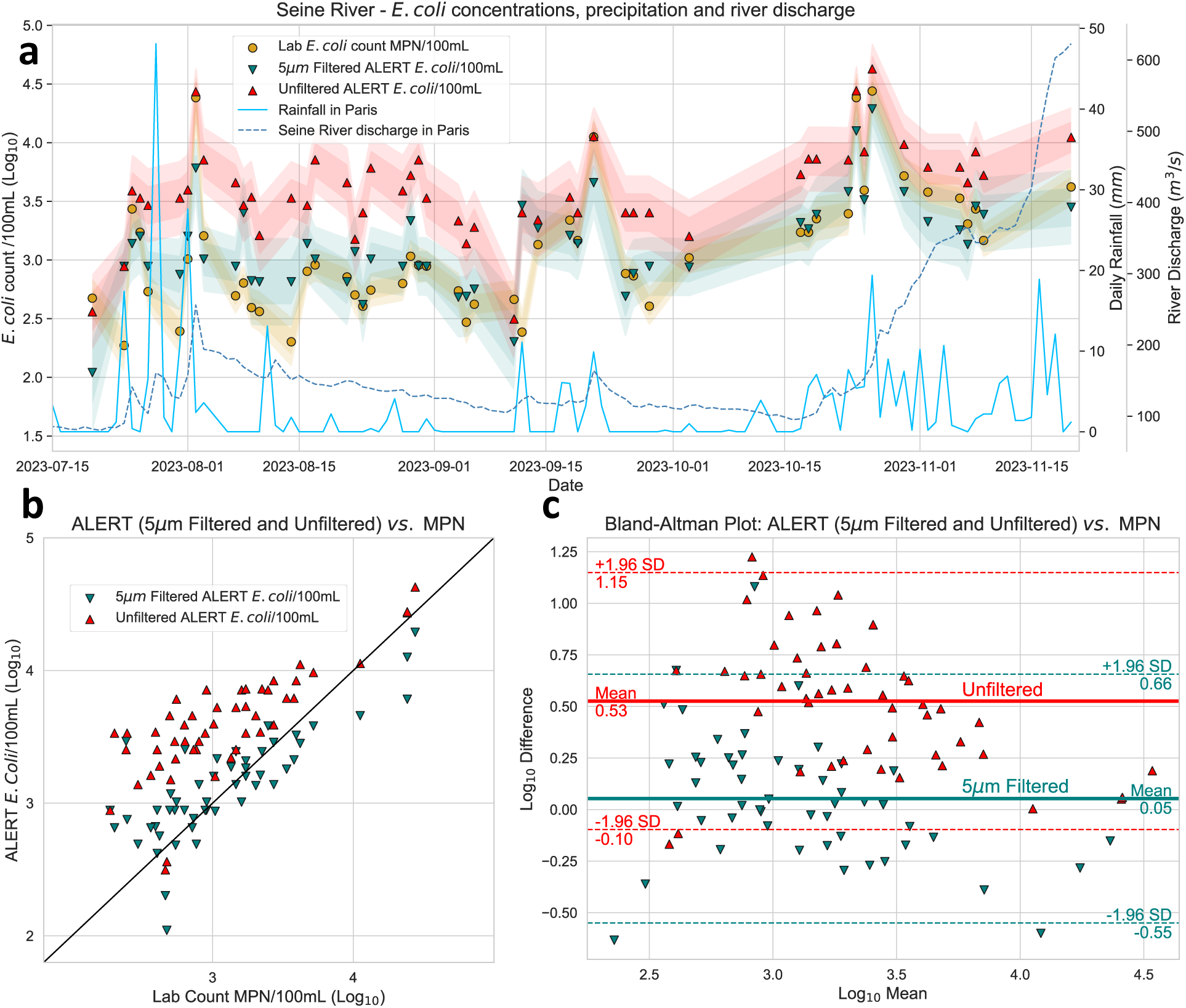
**a**: E. coli concentration time series in the Seine River at the 2024 Olympic venue, obtained via ALERT for both 5 µm-filtered (blue triangles) and unfiltered samples (red triangles), juxtaposed with MPN data (orange circles) and local environmental factors such as daily precipitation (light blue line) and mean river discharge (dashed blue line). Standard deviation and 95% CI shown as colored bands. **B**: Side-by-side analysis of E. coli ALERT measurements for samples filtered at 5 µm and unfiltered, against MPN results. **C**: Same data represented as superposed Bland-Altman plots (mean shown as solid line, 95% CI as dashed lines). While 5 µm-filtered ALERT measurements provide good overall agreement with MPN measurements (0.05 Log_10_ units mean difference, corresponding to a factor of 1.1), consistently higher counts are obtained from unfiltered ALERT measurements, surpassing MPN estimates by an average of 0.53 Log_10_ units, corresponding to a factor of 3.4, thus highlighting the substantial presence of E. coli bound to larger aggregates.

Comparative analyses using side-by-side and Bland-Altman plots (Figures 1b,c) showed good agreement between 5 µm-filtered ALERT and MPN *E. coli* measurements according to the USEPA site-specific criteria (index of agreement IA=0.84)^52^, with a slight mean Log_10_ difference of +0.05 (12% higher) within overlapping 95% confidence intervals. In unfiltered samples, on the other hand, the ALERT method detected significantly higher *E. coli* concentrations compared to MPN, with a mean difference of +0.53 Log_10_ units (239% increase) and high variability: differences as high as +1.22 Log_10_ units (1576% increase) have been recorded. This discrepancy, confirmed by non-overlapping 95% confidence intervals and distinct groupings in the comparative plots, implies a significant yet highly variable FIB load within aggregates larger than 5 µm.

### 3.2 Risk assessment discrepancies due to aggregate-bound FIB

It is instructive to compare the MPN FIB counts with ALERT whole-sample comprehensive counts measured in raw (unfiltered) samples, in terms of the statistical threshold value (STV) and the geometric mean value (GM), regulatory risk assessment parameters whose repeated exceedances may trigger actions such as temporary or permanent closures of a recreational site. Since regulatory GM and STV values differ significantly between the United States (GM=126, STV=410)^28^ and Europe (GM=900 (90^th^ percentile), STV=1800), agreement matrices were computed for both standards (Table 1). According to the stricter U.S. standards, in 100% (GM) and, respectively, 78.4% (STV) of the cases both methods agreed that water quality was insufficient for recreational activities. According to the European standards, on the other hand, agreement rate was only 58.8% (GM) and, respectively, 43.1% (STV). The discrepancies between the two methods were systematically caused by MPN values registering below, and ALERT values above, the threshold, leading to frequent contradictory risk management recommendations that are attributable to aggregate-bound FIB load.

**Table 1.** Agreement matrices for E. coli concentration results obtained by the MPN and ALERT methods on raw samples, in terms of the STV and GM threshold criteria recommended in the United States (left two panels) and Europe (right two panels)

| United States GM Criterion |  |  |  | United States STV Criterion |  |  |  | European GM Criterion |  |  |  | European STV Criterion* |  |  |  |
| --- | --- | --- | --- | --- | --- | --- | --- | --- | --- | --- | --- | --- | --- | --- | --- |
| t=126 | ALERT<t | ALERT≥t | Total | t=410 | ALERT<t | ALERT≥t | Total | t=900 | ALERT<t | ALERT≥t | Total | t=1800 | ALERT<t | ALERT≥t | Total |
| MPN<t | 0 | 0 | 0 | MPN<t | 0 | 9 | 9 | MPN<t | 3 | 21 | 24 | MPN<t | 7 | 29 | 36 |
| MPN≥t | 0 | 51 | 51 | MPN≥t | 2 | 40 | 42 | MPN≥t | 0 | 27 | 27 | MPN≥t | 0 | 15 | 15 |
| Total | 0 | 51 | 51 | Total | 2 | 49 | 51 | Total | 3 | 48 | 51 | Total | 7 | 44 | 51 |
| Agreement rate 100,0% |  |  |  | Agreement rate 78,4% |  |  |  | Agreement rate 58,8% |  |  |  | Agreement rate 43,1% |  |  |  |

\*Specific countries or sites in Europe may apply different STV criteria

### 3.3 Size fractionation investigation

We employed a size fractionation approach to investigate the prevalence of FIB across aggregates of different sizes, with analyses performed on a series of twelve samples from the Seine River collected between mid-October and mid-November 2023. The fractionation process involved successive filtrations using filters with decreasing pore diameters, specifically 10 µm and 5 µm. Filtrations through pore sizes larger than 10 µm were attempted initially but had to be abandoned due to observed disaggregation of larger entities under filtration stress. This approach allowed free bacteria and aggregates below the filter cutoff to freely flow through while retaining aggregates larger than the cutoff, thus facilitating the estimation of the overall FIB load associated with aggregates of varying sizes and the quantification of FIB load per individual aggregate. In both raw and sequentially filtered samples, the MPN method was used to quantify the number of FIB-containing entities (free bacteria or aggregates) while ALERT was used to measure the total FIB load (Figure 2a-c).

**Figure 2.**
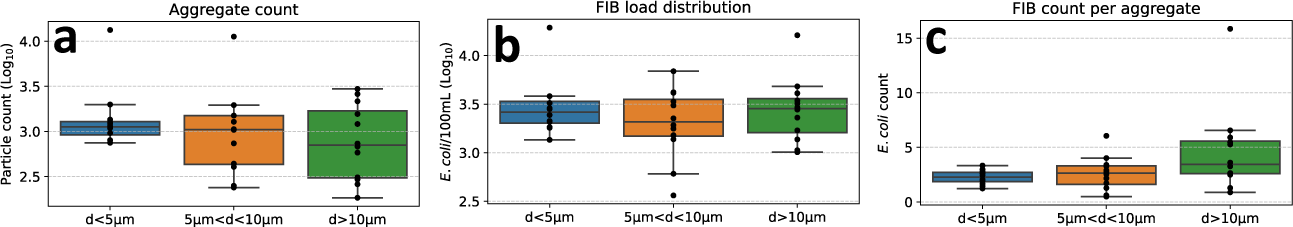
Size fractionation study of aggregates in n=12 Seine River samples by cascade filtration using different pore-size filters. a: Distribution of FIB-containing aggregates across size categories, from differential MPN measurements. b: Distribution of the total FIB (E. coli) load, from differential ALERT measurements. c: Average FIB count distribution per aggregate, from the ratio of the previous two numbers. The box plot center line represents the median, bounds of box the interquartile range (IQR), whiskers extend from box edges to the smallest and largest values within 1.5 times the corresponding IQR.

The count of small entities (free bacteria and aggregates with diameter *d*<5 µm) showed a relatively narrow distribution, with the number of aggregates in higher size categories (5 µm<*d*<10 µm and *d*>10 µm) showing progressively lower values but higher variability. Intriguingly, the total FIB load was, on average, uniformly dispersed among the three size categories, with the most significant fluctuation observed within the 5 µm<d<10 µm range. Consistent with expectations, smaller aggregates exhibited a lower average number of *E. coli* per aggregate, with a mean of 2.3 for entities under 5 µm (maximum observed: 3.3). For larger aggregates, both the average *E. coli* count and its variability increased: aggregates sized between 5 µm and 10 µm harbored an average of 2.6 *E. coli* (maximum: 6.1), while those larger than 10 µm held an average of 4.65 *E. coli* per aggregate (maximum: 15.9).

### 3.4 Microscopic and molecular confirmation of aggregates

Our results were substantiated by microscopic analyses following 0.2 µm filtration and direct on-membrane staining with a range of fluorescent dyes. Figure 3a,b showcases sections of a micrograph from a sample (collected on September 14, 2023), stained with a highly permeable fluorescent dye that labels all bacteria, independent of Gram status. Figure3a reveals dispersed individual bacteria, whereas Figure 3b illustrates a relatively large (30 µm) aggregate with numerous attached bacteria. We subsequently enhanced the staining specificity for Gram-positive bacteria (Figure 3c,d, sample collected on October 30, 2023), discerning aggregates ranging from 10 µm to over 70 µm, with both Gram-positive and Gram-negative bacteria present, suggesting the coexistence of potential enterococci and *E. coli*. The counts of aggregate-bound bacteria align with the size fractionation data depicted in Figure 2c. While more specific staining procedures for *E. coli* exist, they were avoided due to the known risk of disturbing the structure of the aggregates^53^, the integrity of which is central to our study.

**Figure 3.**
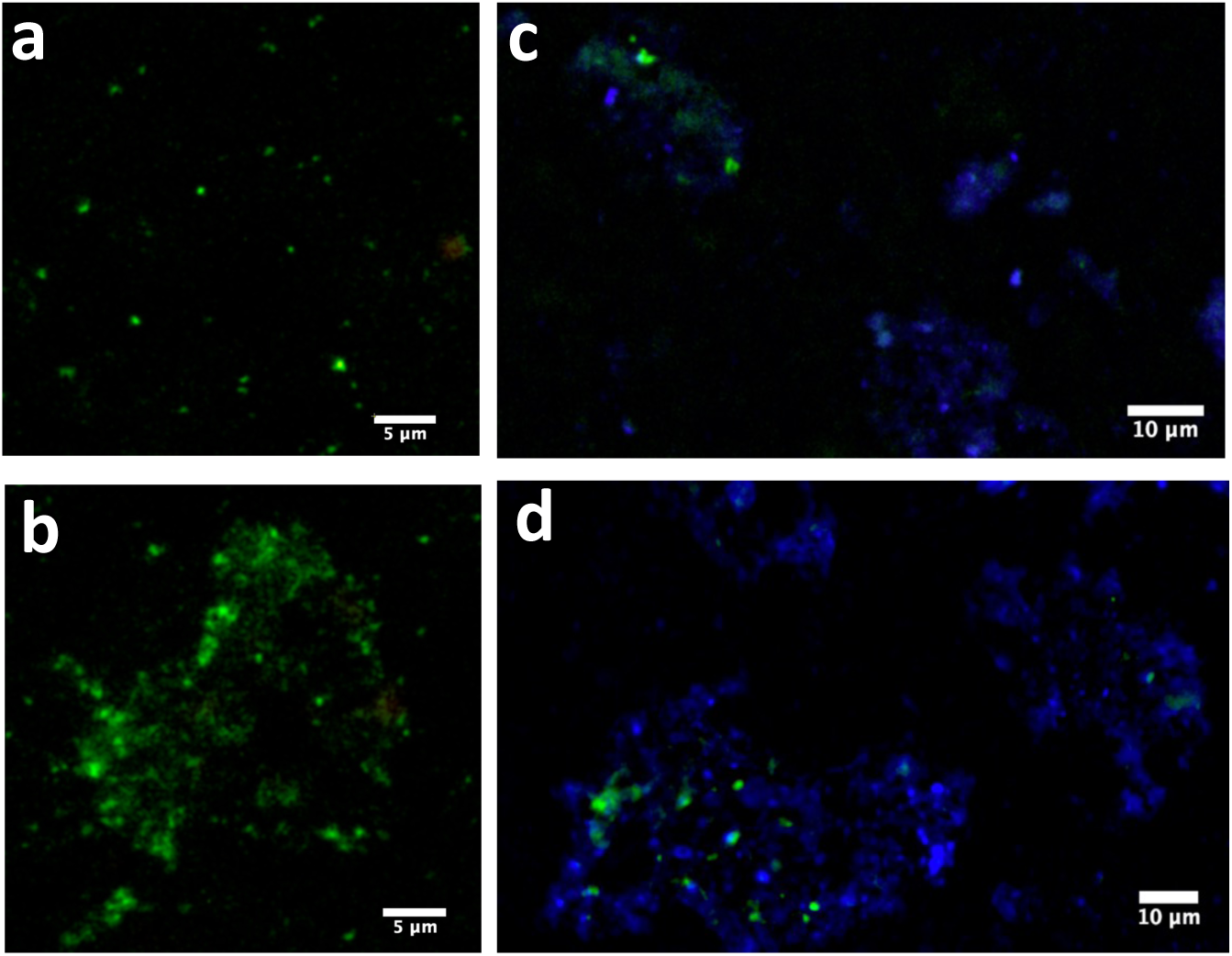
Left: Two portions of the same epi-fluorescence micrograph (Sept. 14, 2023), displaying simultaneous presence of isolated bacteria (a) and aggregates (b). BactoViewTM Live Green staining was applied to visualize all bacteria (green fluorescence). Right: Micrographs (c,d) of bacterial aggregates in a sample collected on Oct. 30, 2023, providing microscopic evidence of the diversity in size and composition of bacterial aggregates. All bacteria were stained with DAPI (blue fluorescence), while gram-positive species were stained with CF488A WGA (green fluorescence), identifying likely candidates for E. coli and Enterococci, respectively.

A striking contrast is observed between the samples: the September sample, collected at the end of the dry summer season, displayed numerous individual bacteria and only occasional aggregates, whereas the October sample, obtained after the onset of the rainy season, was characterized by the prevalence of bacterial aggregates. This trend was corroborated by ALERT assessments of the samples, both unfiltered and after 5 µm filtration. The September sample registered a modest surplus of *E. coli* on aggregates, quantified at +0.07 Log_10_ units (17% surplus), whereas the October sample demonstrated significantly larger surplus of +0.40 Log_10_ units (151% surplus). However, neither precipitation nor river discharge seem to consistently explain the observed abundance of aggregate-bound FIB load: notably, some of the most substantial loads were detected during the driest interval from mid-August to early September, while markedly different aggregate-bound fractions were recorded during the low river discharge conditions in mid-September and mid-October. Such inconsistencies could be explained by combined sewer infrastructure dysfunctions.

Molecular RT-qPCR analyses in three samples collected at the end of the 2023 dry season (September 25, 26, 28) similarly confirmed a significant surplus of *E.coli* RNA in unfiltered compared to 5 µm-filtered samples (122%, 70% and, respectively, 144% surplus), coherent with ALERT measurements (417%, 231%, 187% surplus), and exceeding the planktonic counts often by substantial multiples. By contrast, the MPN method proved insensitive to aggregate-bound FIB, displaying only minimal variation (20%, 16% and, respectively, 7% surplus) between unfiltered and filtered samples (Figure 5e).

### 3.5 Prevalence of aggregate-bound FIB in other water matrices

To understand whether the presence of aggregate-bound *E. coli* bacteria in the Seine river was a localized or widespread phenomenon we applied similar methodology to evaluate FIB surplus in unfiltered versus 5 µm-filtered samples from several water matrices across multiple geographical locations: irrigation waters in California’s Central Valley, treated wastewater effluent from a facility in Milan, Italy, the initial stormwater runoff in the Los Angeles River, USA, and coastal waters during a swimming competition in Brighton, UK.

Figure 4a highlights pronounced differences in irrigation canal samples, with culturable *E. coli* associated with aggregates surpassing planktonic forms by up to two orders of magnitude, a phenomenon likely attributable to sediment resuspension in flowing canals. Conversely, reservoir samples, benefiting from prolonged sedimentation, exhibited minimal aggregate presence, despite similar overall FIB concentrations.

**Figure 4.**
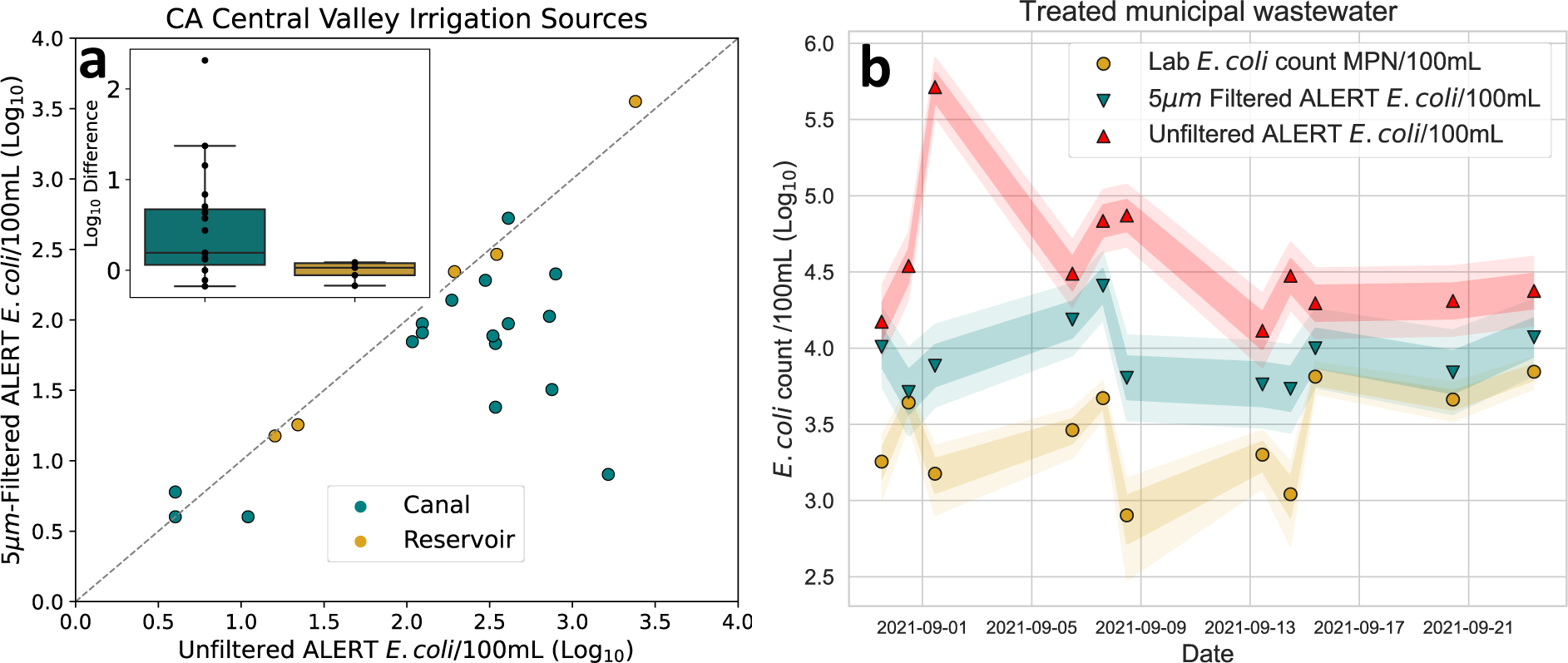
Comparative E. coli quantification in irrigation water sources (left) and treated wastewater (right). a: E. coli concentration disparity in California’s Central Valley irrigation canals (blue, n=19) and reservoirs (gold, n=5), contrasting 5 µm-filtered (vertical axis) against unfiltered (horizontal axis) samples. The box plot of the Log10 difference between unfiltered and filtered values (inset - center line represents the median, bounds of box the IQR, whiskers extend from box edges to the smallest and largest values within 1.5 times the corresponding IQR) reveals high variability and important E. coli load on aggregates for canal samples — up to 100-fold in unfiltered compared to 5 µm-filtered samples—attributed to sediment disturbance. Reservoirs, allowing ample time for sedimentation, show minimal aggregation effects. b: Time series of E. coli concentrations in treated municipal wastewater designated for crop irrigation, with standard deviations and 95% CI shown as colored bands. The important aggregate-associated E. coli loads visible from the ALERT data are not detected by standard MF techniques, leading to occasional 100-fold and higher disparities.

**Figure 5.**
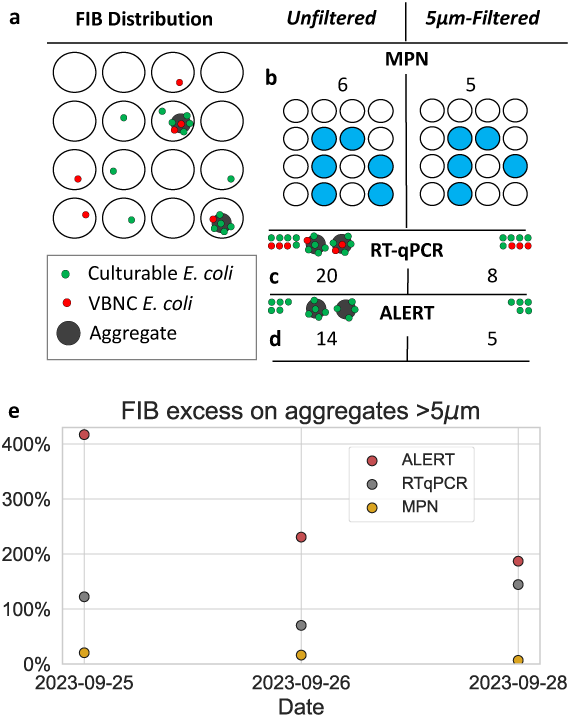
a: Schematic of hypothetical FIB distribution among free and aggregate-bound states, demonstrating (b) that the MPN method can be insensitive to the aggregate-bound FIB load, while RT-qPCR and ALERT can detect substantial presence of bound viable and, respectively, culturable FIB (the difference between RT-qPCR (c) and ALERT (d) being attributed to VBNC bacteria). e: The analysis of relative FIB excess observed on aggregates larger than 5 µm, measured on three distinct Seine River samples using MPN, RT-qPCR and ALERT. The September 28, 2023, results (MPN: 7%, RT-qPCR: 144%, ALERT: 187%) resemble those of a hypothetical distribution schematized in Panel a (20%, 150%, 180%).

Figure 4b captures a month-long time series of MF, 5 µm-filtered, and unfiltered ALERT *E. coli* counts in effluent samples designated for agricultural reuse, from a municipal wastewater treatment facility (collected pre-UV disinfection). The ALERT unfiltered method consistently registered significantly higher *E. coli* counts compared to the MF method, a systematic phenomenon evidenced by distinct confidence intervals and disparities sometimes exceeding hundredfold, underscoring the MF technique’s inability to assess the total FIB load. Concurrently, a notable quantity of FIB was detected even on aggregates smaller than 5 µm. During the initial stages of the monitoring period, the differences among the three analytical methods were very pronounced. However, towards the latter part of the deployment, the convergence among MF, 5 µm-filtered, and unfiltered ALERT measurements improved, implying a possible transient anomaly in the treatment facility’s operations. This was substantiated by independent assessments of high suspended solids levels in the treated effluent (not shown).

Stormwater samples from the Los Angeles River, collected during the first major rainfall of the 2023 wet season on November 16, revealed similar significant discrepancies: duplicate analyses using the ALERT method for unfiltered samples produced a geometric mean *E. coli* concentration of 5.6×10^4^ per 100mL, whereas samples filtered at 5 µm exhibited a mean of only 2.7×10^4^ *E. coli*/100mL. A possible explanation is that, even when the presence of an extensive municipal separate storm sewer system prevents CSOs, traditional standard methods might not capture a considerable fraction of the FIB load during sudden hydrological events.

In samples collected from various locations along the southern coast of Britain in 2023, notable variances between filtered and unfiltered samples were detected, hinting at the presence of FIB aggregates even in instances of very low, or undetectable, planktonic *E. coli* levels. Such variances could result in conflicting risk assessments. For instance, data from a swimming competition in Brighton, UK, in September 2023, showed no detectable FIB in 5 µm-filtered samples but provided unfiltered sample results of 575 *E. coli*/100mL, suggesting unacceptable associated health risk. Such significant variability in comprehensive FIB measurements, also observed in molecular assays^49^, is bound to occur when aggregates, potentially embedded within sediment and shielded from environmental factors such as UV radiation, are dislodged and resuspended in otherwise clean water, especially under the turbulent conditions frequently encountered in near-shore coastal waters.

## 4. Discussion

Our survey across various natural, urban and wastewater environments in four different countries has confirmed the quasi-ubiquitous presence of aggregate-bound FIB, leading to very significant and highly variable ratios of total FIB count to that from traditional MPN or MF methods. Despite reported evidence of aggregate presence^46,47,48^ and their critical implications for risk assessment practices, aggregate-bound FIB and pathogens have been systematically overlooked by existing global regulations, a shortfall attributable to the limitations of conventional MPN and MF quantification methodologies. In the MPN method, the sample is separated into multiple discrete wells which undergo individual presence/absence tests, the distribution of positive reactions allowing determination of the MPN value using probability tables. The MPN method is based on the fundamental assumption that clumps or aggregates do not occur, or else it is prone to underestimating the true microbial density^29^. Indeed, the presence of culturable FIB in planktonic or in aggregate-bound state will produce an identical positive reaction irrespective of whether the corresponding well harbors a single or multiple aggregated bacteria. This results in underrepresentation of the total FIB count when aggregates are present. This limitation extends to earlier variants such as multiple tube fermentation, but also to the MF technique, which similarly fails to differentiate individual bacteria from aggregates in colony counts^28^.

Conversely, molecular assays apply a comprehensive approach, beginning with sample homogenization to release nucleic acids from all cells (planktonic or aggregate-bound), followed by PCR amplification. These assays are sensitive to aggregate-bound FIB but do not discern between culturable and viable but non-culturable (VBNC) cells. They also have different responses to cell viability, with RT-qPCR variants offering enhanced specificity for viable cells^33^.

The ALERT methodology used in this research^51^ merges the culture-based features of the MPN assay with the comprehensive assessment capabilities and detection mechanisms of molecular protocols. Like MPN methods, the ALERT protocol involves mixing the sample with a target-specific combination of growth medium and enzymatic substrate. However, unlike MPN, ALERT does not separate the sample into individual aliquots, but automatically incubates the entire sample while performing continuous monitoring of the ongoing enzymatic reactions via real-time readings of fluorescence and absorbance at specific wavelength combinations. Precise regulation of incubation temperature is critical in the ALERT protocol and is provided by the automated ALERT instrumentation, allowing control of the bacterial lag phase and ensuring consistent subsequent division cycles, which results in a linear correlation between time and the number of bacterial replication cycles. The duration required for the enzymatic signal to be detected and exceed a predetermined threshold thus becomes linearly related to the logarithm of the initial count of target FIB within the sample, which is calculated using the device’s calibration for the specific water matrix (*e.g.* freshwater, seawater, wastewater). Consequently, ALERT’s quantification paradigm is closely related to that of molecular methods, with regular cellular division replacing the DNA amplification in PCR thermal cycles, and the enzymatic signal detection time replacing the cycle threshold Ct.

Traditional MPN and MF culture-based techniques, molecular assays, and ALERT exhibit differential sensitivity not only to aggregate-bound FIB, but also to the physiological state of the bacteria: culturable, VBNC, or nonviable. ALERT can comprehensively and automatically quantify culturable FIB (including aggregate-bound forms) with no interference from VBNC or non-viable bacteria, therefore providing capabilities that are complementary to current molecular methods, while better aligned with global regulatory paradigms which currently rely on viable FIB. As schematically illustrated by the hypothetical FIB distribution shown in Figure 5a, the differential sensitivity of MPN, molecular, and ALERT techniques to FIB aggregates and physiological state can generate significantly different FIB quantification results (Figure 5b-d). Figure 5e provides empirical support for this conceptual framework, presenting data from three distinct Seine River samples analyzed by MPN, RT-qPCR, and ALERT.

Previous studies on bacterial aggregates have predominantly concentrated on spontaneous assembly, resulting in their universal presence in natural environments^44,45,46,54^. Yet, aggregate-bound FIB and associated pathogens, critical from a public health perspective, typically do not emerge from ad-hoc aggregation in the water column. Instead, they originate from the systematic breakdown of fecal matter, where they are embedded within a complex low-density organic matrix^55^. In the microscopic size ranges reported here, aggregates resist further fragmentation under typical hydrodynamic forces, while their buoyancy, aided by Brownian motion and turbulence, ensures their persistence in the water column. These structures not only provide a protective chemical microenvironment^45^ that enhances the resilience of FIB and pathogens to environmental stressors^56^ and typical disinfection processes used in water treatment^47^, but can also increase their infectivity by providing immune evasion mechanisms^45^ and antibiotic resistance^46^. Frequently, aggregates are released into environmental waters without treatment, through phenomena such as CSOs or stormwater influx, or they may infiltrate from agricultural runoff or septic systems into potable water resources. Our study shows that their substantial load of FIB and pathogens, directly derived from the originating fecal matter, is detected in most water matrices included in the study yet escapes detection by the prevalent FIB quantification techniques.

The measured ratio of total FIB count to that from traditional MPN or MF methods is not only substantial but also highly variable, ranging from an average of 3.4 measured in the Seine River over a bathing season, to over two orders of magnitude recorded on specific samples from irrigation canals and treated municipal wastewater effluent. Due to their significant variability, such effects cannot be accounted for by applying a simple “renormalization” or corrective factor to existing risk limits. Our study shows that they have major implications for risk assessment practices, leading to frequent contradictory assessments against regulatory concentration thresholds and significant underestimation of risk levels by traditional MPN/MF culture-based techniques (Table 1). These results could explain the ongoing prevalence of public health issues associated with contaminated drinking water^1^, irrigation practices^4^, and open-water recreational and athletic activities^6^, which continue to arise despite intensified regulatory surveillance. They also offer insight into the observed discrepancies between qPCR and culture-based methods, providing a potential explanation for the extreme variabilities in FIB measurements frequently reported in recreational^49^ and stormwater^50^ samples.

## 5. Conclusions

Our research demonstrates that the existing limitations of current regulatory frameworks based on traditional MPN and MF techniques render them inadequate for assessing the microbiological risks posed by aggregate-bound FIB and associated pathogens in water, despite their high abundance across different water matrices. The integration of comprehensive FIB analysis techniques, including established molecular methods in combination with rapid culture-based approaches like those used in the present study, is essential to overcome these limitations. Such techniques provide water safety assessments of highly improved accuracy, with critical implications for new epidemiological investigations and for designing improved regulatory frameworks. The adoption of automated field technologies that can provide comprehensive quantification of culturable fecal indicators could help combat the critical public health impact of microbiologically unsafe water, enabling risk monitoring including in remote and resource-limited settings. The methodology developed in the present study represents a promising advance toward resolving these critical public health challenges.

## Data availability

Data that support the findings of this study are available within the paper or upon request. Historical rainfall in Paris (measured at the Montsouris station) was obtained from: https://prevision-meteo.ch/climat/mensuel/paris-montsouris. Historical Seine River discharge data (measured at the Austerlitz bridge in Paris) was obtained from: https://www.hydro.eaufrance.fr/stationhydro/F700000103/fiche

## Code availability

All graphs were generated in Python using the matplotlib and seaborn libraries. All scripts and data available upon request. Image processing was performed using the Fiji distribution of the ImageJ software, available from:

https://imagej.net/software/fiji/downloads

## Inclusion and Ethics

No ethical considerations apply to this research.

## Acknowledgements

Figure 4a data resulted from research supported by NIFA/USDA SBIR grant 2019-33610-29766, with fieldwork acknowledgments to Dr. Trevor Suslow’s team (U.C. Davis). Figure 4b data were obtained through research supported by the Digital Water City project, with funding from the European Union’s H2020 Research and Innovation Programme under Grant Agreement No. 820954, and acknowledgements for installation support and laboratory contributions to Marco Bernardi and the Gruppo CAP team. We acknowledge Omar Bach-Rais, Kai Hong, Patrick Ea, Maelys Hemon, Andreas Hausot and Vaizanne Huynh for technical support contributions on different parts of the project. We thank Mark Cooper for performing sampling and ALERT analyses at the Brighton, UK swimming competition. We are grateful to Ion Alexandru Bobulescu, Victoire Rerolle, Michelle Yeh, and Nic Volanschi for fruitful discussions and critical review.

## Author contributions

DEA conceived the project, coordinated European activities, developed the laboratory and imaging protocols, performed image processing and data analysis and interpretation. He also drafted the manuscript. DAS performed instrument maintenance, upgrades, and coordinated sampling campaigns. RG performed daily sampling from the Seine River, sample filtrations, laboratory microbiology work, and much of the microscopy work. DW performed stormwater sampling from the Los Angeles River, and associated microbiology work. JW oversaw the U.S. project activities and coordinated the agricultural sampling and data analysis. All authors reviewed the manuscript in detail.

## References

1 World Health Organization. State of the World’s Drinking Water: an urgent call to action to accelerate progress on ensuring safe drinking water for all. ISBN 978-92-4-006080-7 (2022).

2 UNICEF. Triple Threat: How disease, climate risks, and unsafe water, sanitation and hygiene create a deadly combination for children. ISBN: 978-92-806-5438-7 (2023).

3 World Health Organization. Guidelines for Drinking Water Quality, Fourth Edition incorporating the first and second addenda. ISBN 978-92-4-004506-4 (2022).

4 Marshall, K. E. et al. Lessons Learned from a Decade of Investigations of Shiga Toxin–Producing *Escherichia coli* Outbreaks Linked to Leafy Greens, United States and Canada. Emerg. Infect. Dis. 26, 2319–2328 (2020).

5 Spalding, A., Goodhue, R. E., Kiesel, K. & Sexton, R. J. Economic impacts of food safety incidents in a modern supply chain: *E. coli* in the romaine lettuce industry. Am. J. Agric. Econ. 105, 597–623 (2023).

6 DeFlorio-Barker, S., Wing, C., Jones, R. M. & Dorevitch, S. Estimate of incidence and cost of recreational waterborne illness on United States surface waters. Environ. Heal. 17, 3 (2018).

7 Intergovernmental Panel on Climate Change. Climate change 2022: Impacts, adaptation, and vulnerability. Working Group II Contribution of to the Sixth Assessment Report of the Inter-governmental Panel on Climate Change. (2021).

8 Drechsel, P., Qadir, M. & Galibourg, D. The WHO Guidelines for Safe Wastewater Use in Agriculture: A Review of Implementation Challenges and Possible Solutions in the Global South. Water 14, 864 (2022).

9 Overbury, K., Conroy, B. W. & Marks, E. Swimming in nature: A scoping review of the mental health and wellbeing benefits of open water swimming. J. Environ. Psychol. 90, 102073 (2023).

10 Associated Press: “Water quality worries force cancelation of Paris Olympics swimming test event in Seine River”, https://apnews.com/article/paris-olympics-swimming-test-seine-river-canceled-ee1541c92d188de458ea479cf9639a0c (2023)

11 BBC: “Dozens fall ill after Sunderland triathlon, health chiefs confirm” https://www.bbc.com/news/uk-england-tyne-66421422 (2023)

12 Cabelli, V. J. Health Effects Criteria for Marine Recreational Waters, EPA 600/1-80-031, U.S. Environmental Protection Agency (1983).

13 Dufour, A. P. Health Effects Criteria for Fresh Recreational Waters, EPA-600/1-84-004, U.S. Environmental Protection Agency (1984).

14 Kay, D. et al. Derivation of numerical values for the World Health Organization guidelines for recreational waters. Water Res. 38, 1296–1304 (2004).

15 UNICEF. Target Product Profile: Rapid *E*. coli Detection Tests (2019).

16 World Health Organization. Progress on household drinking water, sanitation and hygiene 2000-2020: five years into the SDGs. ISBN 978-92-4-003084-8 (2021).

17 International Organization for Standardization. ISO 5667-1 Water quality — Sampling — Part 1: Guidance on the design of sampling programmes and sampling techniques (2020).

18 International Organization for Standardization. ISO 5667-2 Water quality — Sampling — Part 2: Guidance on sampling techniques (1991).

19 International Organization for Standardization. ISO 5667-3 Water quality — Sampling — Part 3: Preservation and handling of water samples (2018).

20 International Organization for Standardization. ISO 9308-1 Water quality — Enumeration of Escherichia coli and coliform bacteria - Part 1: Membrane filtration method for waters with low bacterial background flora (2014).

21 International Organization for Standardization. ISO 7899-2 Water quality — Detection and enumeration of intestinal enterococci — Part 2: Membrane filtration method. (2000).

22 U.S. Environmental Protection Agency. EPA Method 1603: Escherichia coli (E. coli) in Water by Membrane Filtration Using Modified membrane-Thermotolerant Escherichia coli Agar (Modified mTEC) (2002).

23 U.S. Environmental Protection Agency. EPA Method 1600: Enterococci in Water by Membrane Filtration Using membrane-Enterococcus Indoxyl-Beta-D-Glucoside Agar (mEI) (2009).

24 International Organization for Standardization. ISO 9308-2 Water quality — Enumeration of Escherichia coli and coliform bacteria Part 2: Most probable number method (2012).

25 International Organization for Standardization. ISO 9308-3 Water quality — Enumeration of Escherichia coli and coliform bacteria Part 3: Miniaturized method (Most Probable Number) for the detection and enumeration of E. coli in surface and waste water (1998).

26 International Organization for Standardization. ISO 7899-1 Water quality — Detection and enumeration of intestinal enterococci in surface and waste water —Part 1: Miniaturized method (Most Probable Number) by inoculation in liquid medium. (1998).

27 American Society for Testing and Materials. ASTM D6503-19 Standard Test Method for Enterococci in Water Using Enterolert. (2019)

28 U.S. Environmental Protection Agency. Recreational Water Quality Criteria, USEPA Report 820-F-12-058 (2012).

29 Jarvis, B., Wilrich, C. & Wilrich, P. -T. Reconsideration of the derivation of Most Probable Numbers, their standard deviations, confidence bounds and rarity values. J. Appl. Microbiol. 109, 1660–1667 (2010).

30 Baird, R. B., Eaton, A. D. & Rice, E. W. Standard Methods for the Examination of Water and Wastewater, 23rd Edition. (2017).

31 Garcia-Armisen, T. & Servais, P. Partitioning and fate of particle-associated *E. coli* in river waters. Water Environ. Res. 81, 21–28 (2009).

32 Passerat, J., Ouattara, N. K., Mouchel, J.-M., Rocher, V. & Servais, P. Impact of an intense combined sewer overflow event on the microbiological water quality of the Seine River. Water Res. 45, 893–903 (2011).

33 Bergeron, P., Oujati, H., Cuenca, V. C., Mestre, J. M. H. & Courtois, S. Rapid monitoring of *Escherichia coli* and Enterococcus spp. in bathing water using Reverse Transcription-quantitative PCR. Int. J. Hyg. Environ. Heal. 214, 478–484 (2011).

34 Nocker, A., Cheung, C.-Y. & Camper, A. K. Comparison of propidium monoazide with ethidium monoazide for differentiation of live vs. dead bacteria by selective removal of DNA from dead cells. J. Microbiol. Methods 67, 310–320 (2006).

35 Yuan, Y., Zheng, G., Lin, M. & Mustapha, A. Detection of viable *Escherichia coli* in environmental water using combined propidium monoazide staining and quantitative PCR. Water Res. 145, 398–407 (2018).

36 U.S. Environmental Protection Agency. Method 1611: Enterococci in Water by TaqMan® Quantitative Polymerase Chain Reaction (qPCR) Assay (2012).

37 U.S. Environmental Protection Agency. Method 1609.1: Enterococci in Water by TaqMan® Quantitative Polymerase Chain Reaction (qPCR) with Internal Amplification Control (IAC) Assay (2015).

38 Botes, M., Kwaadsteniet, M. de & Cloete, T. E. Application of quantitative PCR for the detection of microorganisms in water. Anal. Bioanal. Chem. 405, 91–108 (2013).

39 Wade, T. J. et al. Rapidly Measured Indicators of Recreational Water Quality Are Predictive of Swimming-Associated Gastrointestinal Illness. Environ. Heal. Perspect. 114, 24–28 (2006).

40 Wade, T. J. et al. Health risks to children from exposure to fecally-contaminated recreational water. PLoS ONE 17, e0266749 (2022).

41 Report on the 2nd Five-Year Review of EPA’s Recreational Water Quality Criteria, USEPA Report 822-R-23-003, U.S. Environmental Protection Agency, Office of Water (2023).

42 Wade, T. J. et al. Rapidly measured indicators of recreational water quality and swimming-associated illness at marine beaches: a prospective cohort study. Environ. Heal. 9, 66 (2010).

43 Whitman, R. L. et al. Relationship and Variation of qPCR and Culturable Enterococci Estimates in Ambient Surface Waters Are Predictable. Environ. Sci. Technol. 44, 5049–5054 (2010).

44 Flemming, H.-C. & Wuertz, S. Bacteria and archaea on Earth and their abundance in biofilms. Nat. Rev. Microbiol. 17, 247–260 (2019).

45 Kragh, K. N., Tolker-Nielsen, T. & Lichtenberg, M. The non-attached biofilm aggregate. *Commun*. Biol. 6, 898 (2023).

46 Cai, Y.-M. Non-surface Attached Bacterial Aggregates: A Ubiquitous Third Lifestyle. Front. Microbiol. 11, 557035 (2020).

47 Chahal, C. et al. Pathogen and Particle Associations in Wastewater: Significance and Implications for Treatment and Disinfection Processes (Chapter 2). Adv. Appl. Microbiol. 97, 63–119 (2016).

48 Droppo, I. G. et al. Dynamic Existence of Waterborne Pathogens within River Sediment Compartments. Implications for Water Quality Regulatory Affairs. Environ. Sci. Technol. 43, 1737–1743 (2009).

49 Boehm, A. B. Enterococci Concentrations in Diverse Coastal Environments Exhibit Extreme Variability. Environ. Sci. Technol. 41, 8227–8232 (2007).

50 McCarthy, D. T., Deletic, A., Mitchell, V. G., Fletcher, T. D. & Diaper, C. Uncertainties in stormwater *E. coli* levels. Water Res. 42, 1812–1824 (2008).

51 Angelescu, D. E. et al. Autonomous system for rapid field quantification of *Escherichia coli* in surface waters. J. Appl. Microbiol. 126, 332–343 (2019).

52 U.S. Environmental Protection Agency. Site-specific alternative recreational criteria technical support materials for alternative indicators and methods. EPA-820-R-14-011 (2014)

53 Örmeci, B. & Linden, K. G. Development of a fluorescence in situ hybridization protocol for the identification of micro-organisms associated with wastewater particles and flocs. J. Environ. Sci. Heal., Part A 43, 1484–1488 (2008).

54 Balzer, M., Witt, N., Flemming, H.-C. & Wingender, J. Faecal indicator bacteria in river biofilms. Water Sci. Technol. 61, 1105–1111 (2010).

55 Rose, C., Parker, A., Jefferson, B. & Cartmell, E. The Characterization of Feces and Urine: A Review of the Literature to Inform Advanced Treatment Technology. Crit. Rev. Environ. Sci. Technol. 45, 1827–1879 (2015).

56 Characklis, G. W. et al. Microbial partitioning to settleable particles in stormwater. Water Res. 39, 1773–1782 (2005).

